# A single intranasal dose of chimpanzee adenovirus-vectored vaccine protects against SARS-CoV-2 infection in rhesus macaques

**DOI:** 10.1101/2021.01.26.428251

**Authors:** Ahmed O. Hassan, Friederike Feldmann, Haiyan Zhao, David T. Curiel, Atsushi Okumura, Tsing-Lee Tang-Huau, James Brett Case, Kimberly Meade-White, Julie Callison, Jamie Lovaglio, Patrick W. Hanley, Dana P. Scott, Daved H. Fremont, Heinz Feldmann, Michael S. Diamond

**Affiliations:** Department of Medicine, Washington University School of Medicine, St. Louis, MO 63110, USA; Rocky Mountain Veterinary Branch Division of Intramural Research, NIAID, NIH, Rocky Mountain Laboratories, Hamilton, MT, USA; Department of Pathology & Immunology, Washington University School of Medicine, St. Louis, MO 63110, USA; Department of Radiation Oncology, Washington University School of Medicine, St. Louis, MO 63110, USA; Laboratory of Virology, Division of Intramural Research, NIAID, NIH, Rocky Mountain Laboratories, Hamilton, MT, USA; Department of Biochemistry and Molecular Biophysics, Washington University School of Medicine, St. Louis, MO 63110, USA; Department of Molecular Microbiology, Washington University School of Medicine, St. Louis, MO 63110, USA

## Abstract

The deployment of a vaccine that limits transmission and disease likely will be required to end the Coronavirus Disease 2019 (COVID-19) pandemic. We recently described the protective activity of an intranasally-administered chimpanzee adenovirus-vectored vaccine encoding a pre-fusion stabilized spike (S) protein (ChAd-SARS-CoV-2-S) in the upper and lower respiratory tract of mice expressing the human angiotensin-converting enzyme 2 (ACE2) receptor. Here, we show the immunogenicity and protective efficacy of this vaccine in non-human primates. Rhesus macaques were immunized with ChAd-Control or ChAd-SARS-CoV-2-S and challenged one month later by combined intranasal and intrabronchial routes with SARS-CoV-2. A single intranasal dose of ChAd-SARS-CoV-2-S induced neutralizing antibodies and T cell responses and limited or prevented infection in the upper and lower respiratory tract after SARS-CoV-2 challenge. As this single intranasal dose vaccine confers protection against SARS-CoV-2 in non-human primates, it is a promising candidate for limiting SARS-CoV-2 infection and transmission in humans.

## INTRODUCTION

Severe acute respiratory syndrome coronavirus 2 (SARS-CoV-2) was first isolated in late 2019 from patients with severe respiratory illness in China (Zhou et al., 2020b). SARS-CoV-2 infection results in a clinical syndrome, Coronavirus Disease 2019 (COVID-19) that can progress to respiratory failure (Guan et al., 2020) and systemic inflammatory disease (Cheung et al., 2020; Mao et al., 2020; Wichmann et al., 2020). The elderly, immunocompromised, and those with co-morbidities (*e.g*., obesity, diabetes, and hypertension) are at greatest risk of death from COVID-19 (Zhou et al., 2020a). Virtually all countries and territories have been affected with more than 97 million infections and 2 million deaths recorded worldwide (https://covid19.who.int/). The rapid expansion and prolonged nature of the COVID-19 pandemic and its accompanying morbidity, mortality, and destabilizing socioeconomic effects have made the development and deployment of a SARS-CoV-2 vaccine an urgent global health priority.

The spike (S) protein of the SARS-CoV-2 virion engages the cell-surface receptor angiotensin-converting enzyme 2 (ACE2) to promote coronavirus entry into human cells (Letko et al., 2020). Because the S protein is critical for viral entry, it has been targeted for vaccine development and therapeutic antibody interventions. SARS-CoV-2 S proteins are cleaved to yield S1 and S2 fragments, followed by further processing of S2 to yield a smaller S2’ protein (Hoffmann et al., 2020). The S1 protein includes the receptor binding domain (RBD) and the S2 protein promotes membrane fusion. The prefusion form of the SARS-CoV-2 S protein (Wrapp et al., 2020), which displays the RBD in an ‘up’ position, is recognized by potently neutralizing monoclonal antibodies (Barnes et al., 2020; Cao et al., 2020b; Pinto et al., 2020; Tortorici et al., 2020; Zost et al., 2020) or protein inhibitors (Cao et al., 2020a).

Multiple academic and industry groups are developing vaccine candidates that target the SARS-CoV-2 S protein (Burton and Walker, 2020) using several platforms including DNA plasmid, lipid nanoparticle encapsulated mRNA, inactivated virion, subunit, and viral-vectored vaccines (Graham, 2020), with most requiring two doses. While several vaccines from different platforms are in advanced clinical trials in humans to evaluate safety and efficacy, all lead candidates, including the Pfizer/BioNTech BNT162b2 and Moderna 1273 mRNA vaccines (Baden et al., 2020; Polack et al., 2020) that were granted emergency use authorization (EUA), are administered by intramuscular injection resulting in robust systemic yet poor mucosal immunity. Thus, questions remain as to the ability of these vaccines to curtail both transmission and severe disease, especially if upper airway infection is not prevented or reduced. Indeed, while many of the intramuscular vaccines prevent SARS-CoV-2-induced pneumonia in non-human primates, they variably protect against upper airway infection and presumably transmission (Mercado et al., 2020; van Doremalen et al., 2020; Wang et al., 2020; Yu et al., 2020).

We recently described a chimpanzee Adenovirus (simian Ad-36)-based SARS-CoV-2 vaccine (ChAd-SARS-CoV-2-S) encoding for the S protein. Intranasal administration of a single dose of ChAd-SARS-CoV-2-S induced robust humoral and cell-mediated immune responses against the S protein and prevented upper and lower airway infection in mice expressing the human ACE2 receptor (Hassan et al., 2020). This vaccine differs from ChAdOx1 nCoV-19, a chimpanzee Ad-23-based SARS-CoV-2 vaccine that is currently under phase 3 evaluation in humans as intramuscular injections (NCT04324606): ChAd-SARS-CoV-2-S is derived from a different ChAd serotype, has further deletions in the backbone to enhance production, and introduces proline mutations to stabilize the S protein into a pre-fusion form (Hassan et al., 2020). Here, as a next step toward clinical development of ChAd-SARS-CoV-2-S, we immunized rhesus macaques (*Macaca mulatta*) with a single intranasal dose of ChAd-Control or ChAd-SARS-CoV-2-S, and then challenged animals one month later with SARS-CoV-2 via the combined intranasal and intrabronchial routes. Immunization with ChAd-SARS-CoV-2 resulted in the development of anti-S, anti-RBD, and neutralizing antibodies as well as T cell responses that prevented or limited infection in nasal swabs, bronchoalveolar lavage fluid, and lung tissues after SARS-CoV-2 challenge. Thus, administration of a single dose of the ChAd-SARS-CoV-2-S vaccine through a non-injection route has the potential to protect at the portal of entry and in distant tissues, which could limit both virus-induced disease and transmission.

## RESULTS

### Immunogenicity analysis

We immunized 12 adult rhesus macaques (RM, Indian origin), aged 3 to 11 years old, with ChAd-Control or ChAd-SARS-CoV-2 (n = 6 each; 3 females, 3 males). RM received a single immunization of 10^11^ viral particles of the ChAd vectors by the intranasal route without adjuvant at day −28 (**Fig 1A**). All animals were pre-screened for the absence of neutralizing antibody responses against the simian AdV vector (**Fig S1**) as well as against SARS-CoV-2. Three weeks after vaccination (day −7), we detected IgG antibodies against the S and RBD proteins by ELISA in all RM immunized with ChAd-SARS-CoV-2 but not ChAd-Control vaccines (**Fig 1B-C**). Neutralizing antibody responses were assessed using an infectious SARS-CoV-2 focus-reduction neutralization test (Case et al., 2020) and were detected in all RMs immunized with ChAd-SARS-CoV-2 (**Fig 1D**). T cell responses specific to SARS-CoV-2 S were assessed by an IFN-γ ELISPOT assay on PBMCs isolated two weeks after vaccination (day −14). All RMs immunized with ChAd-SARS-CoV-2 but not ChAd-Control developed T cell responses to SARS-CoV-2 S (**Fig 1E**).

**Figure 1.**
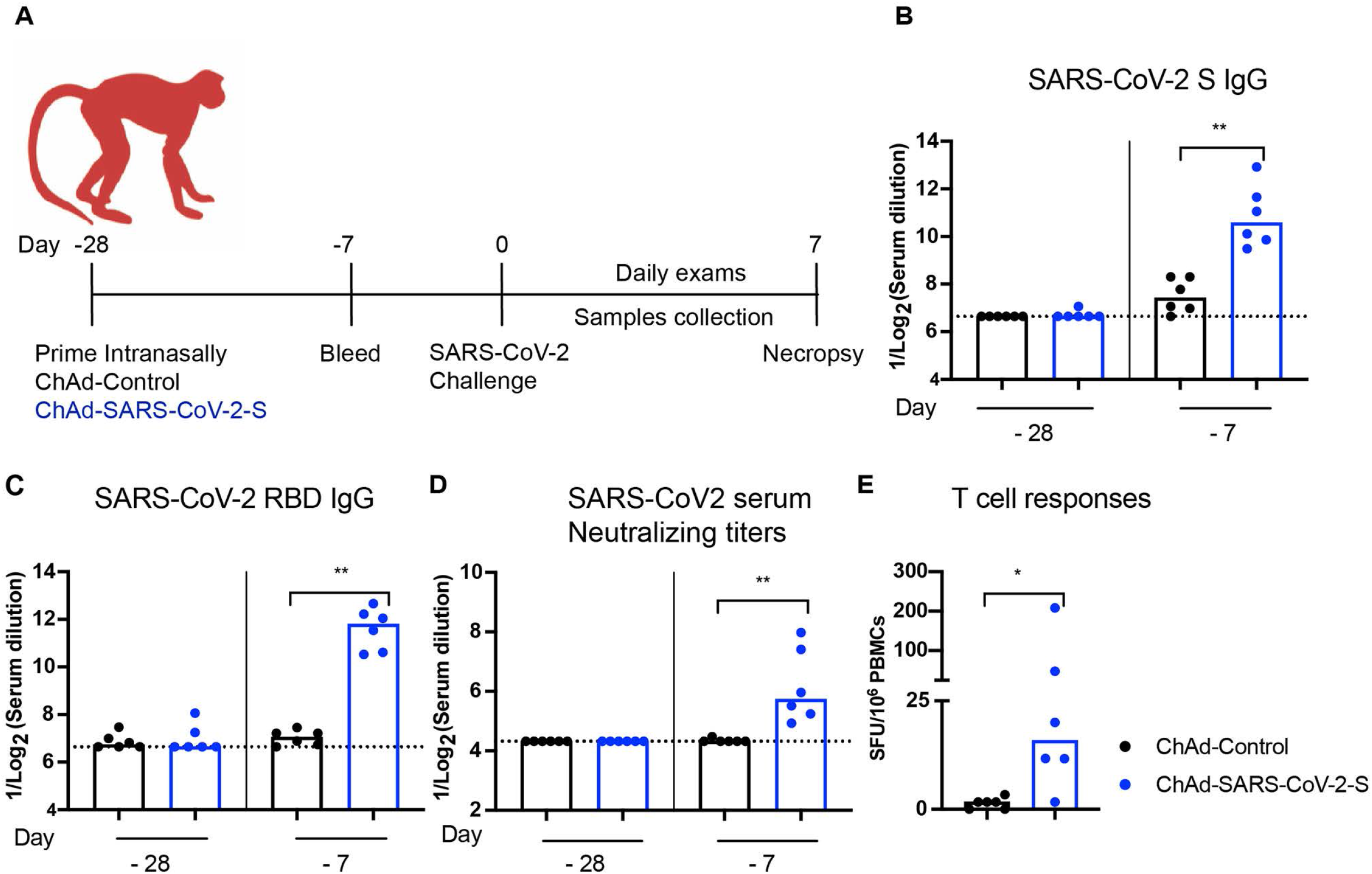
Immunogenicity of ChAd-SARS-CoV-2-S in RM. **A.** RM (3-11 years old) were immunized via intranasal route with a single dose (10^11^ virus particles) of ChAd-Control or ChAd-SARS-CoV-2-S vaccine. Antibody responses in sera of immunized RM at day −28 (day of immunization) and at day −7 (three weeks after immunization) were evaluated. An ELISA measured anti-S and RBD IgG levels (**B-C**), and an FRNT determined neutralization activity (**D**) (n = 6 per group; Mann-Whitney test: **, *P* < 0.01). Bars and columns show median values, and dotted lines indicate the limit of detection (LOD) of the assays. **E**. T cell response. PBMCs were isolated at day −14, and SARS-CoV-2 specific T cells were quantified by counting IFNγ-positive spots following stimulation with a 15-mer SARS-CoV-2 S peptide pool (2 μg/mL). Spot-forming unit counts per 10^6^ stimulated PBMCs after deduction of background counts are shown (n = 6; Mann-Whitney test: *, *P* < 0.05).

### Efficacy of ChAd-SARS-CoV-2 in the upper respiratory tract

At four weeks after immunization, RM were challenged with 1 x 10^6^ fifty-percent tissue culture infective dose (TCID_50_) of SARS-CoV-2 (strain 2019 n-CoV/USA_WA1/2020) split between intranasal and intrabronchial routes of administration. The combined challenge route includes direct infection of the bronchial tree; while not optimal for assaying vaccines that induce mucosal immunity in the upper airway, this protocol was required since intranasal infection, by itself, does not cause consistent lung infection in RMs (Chandrashekar et al., 2020; Muñoz-Fontela et al., 2020). Even though clinical disease is limited in SARS-CoV-2 infected RM (Chandrashekar et al., 2020; Munster et al., 2020) it was less in animals immunized with ChAd-SARS-CoV-2-S than ChAd-Control vaccine (**Fig 2A and Table S1**). Viral loads in the nasal swabs at days +1, +3, +5, and +7 were measured by quantitative reverse transcription PCR (qRT–PCR) using primers for subgenomic (N gene, sgRNA) or genomic (nsp12) RNA. Whereas most of the RMs immunized with ChAd-Control vaccine showed high levels of viral RNA through day+5 with some animals persisting through day +7, lower levels were observed in RMs immunized with ChAd-SARS-CoV-2 (**Fig 2B-E**). At day +1, only one RM vaccinated with ChAd-SARS-CoV-2 showed detectable infectious virus in nasal swabs by a tissue culture infectious dose (TCID_50_) assay, whereas 4 of 6 RM immunized with ChAd-Control were positive (**Fig 2F**). At later time points, infectious virus was not recovered from the nasal swabs of vaccinated animals. Overall, our data suggest that ChAd-SARS-CoV-2 vaccine protects the nasopharyngeal region from SARS-CoV-2 infection and results in reduced viral RNA levels and accelerated clearance.

**Figure 2.**
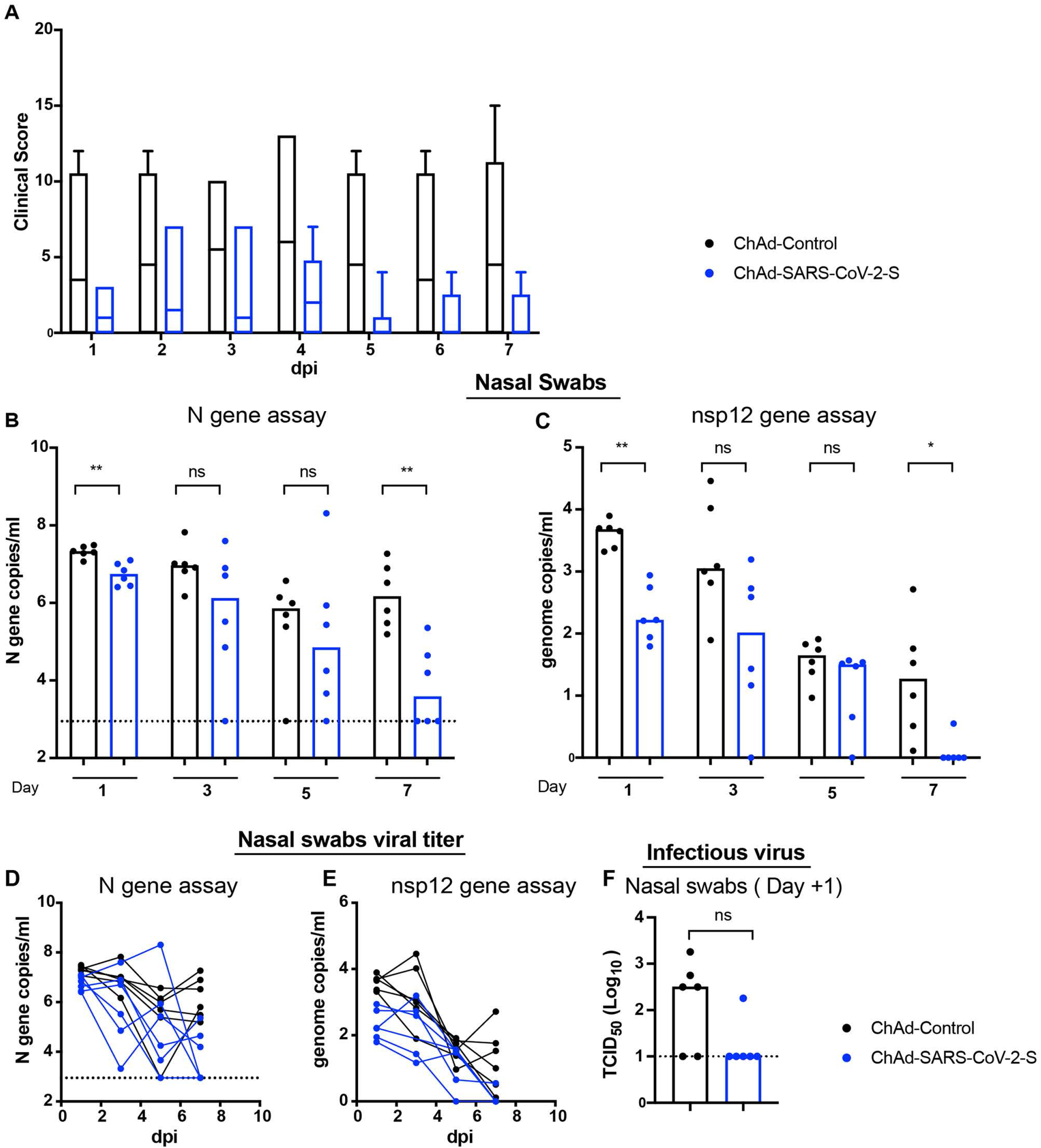
Viral RNA levels in the upper respiratory tract of RM after SARS-CoV-2 challenge. RM (3-11 years old) were immunized via intranasal route with ChAd-Control or ChAd-SARS-CoV-2-S. Four weeks later, RM were challenged with 1×10^6^ fifty-percent tissue culture infective dose (TCID_50_) of SARS-CoV-2 (strain 2019 n-CoV/USA_WA1/2020) split between the intranasal and intrabronchial administration routes. **A.** Clinical scores after challenge. Boxes show the 25th to 75th percentiles, the center lines represent the median, and the extended whiskers from boxes indicate the minimum/maximum values. **B-E.** Nasal swabs were collected at day +1, +3, +5, and +7. Viral RNA was measured by RT–qPCR using primers specific for subgenomic mRNA (N gene) (**B and D**) or genomic RNA (nsp12 gene) (**C and E**) (n = 6, Mann-Whitney test: * *P* < 0.05; ** *P* < 0.01). **D-E**. Longitudinal analysis of SARS-CoV-2 viral RNA in nasal swabs as determined by RT-qPCR showing the subgenomic (**D**) and genomic RNA (**E**). Each connected line shows the viral RNA in individual animal at the indicated time points. **F**. Levels of infectious virus (TCID_50_ analysis) recovered from nasal swabs of RM obtained one day after SARS-CoV-2 challenge (n = 6, Mann-Whitney test: ns, not significant). The dotted line represents the limit of detection of the assay in this Figure.

### Efficacy of ChAd-SARS-CoV-2 in the lower respiratory tract

We next evaluated the protective effects in the lower respiratory tract by measuring infection in bronchoalveolar lavage (BAL) fluid at days +1 and +3 and lung tissues at day +7. For the BAL fluid obtained at day +1, all samples from ChAd-Control immunized RM were positive for infectious virus with titers reaching as high as 10^5^ TCID/ml, whereas only one of six samples from ChAd-SARS-CoV-2-immunized animals was positive with a lower titer of 4 x 10^2^ TCID/ml (**Table 1**). At day +3, only one (from ChAd-control immunized RM) of the 12 BAL fluid samples collected was positive for infectious virus. Consistent with these results, at days +1 and +3, substantially higher levels (100 and 50-fold, respectively) of viral RNA were detected in the BAL fluid of RM immunized with ChAd-control compared to ChAd-SARS-CoV-2 (**Fig 3A-B**).

**Figure 3.**
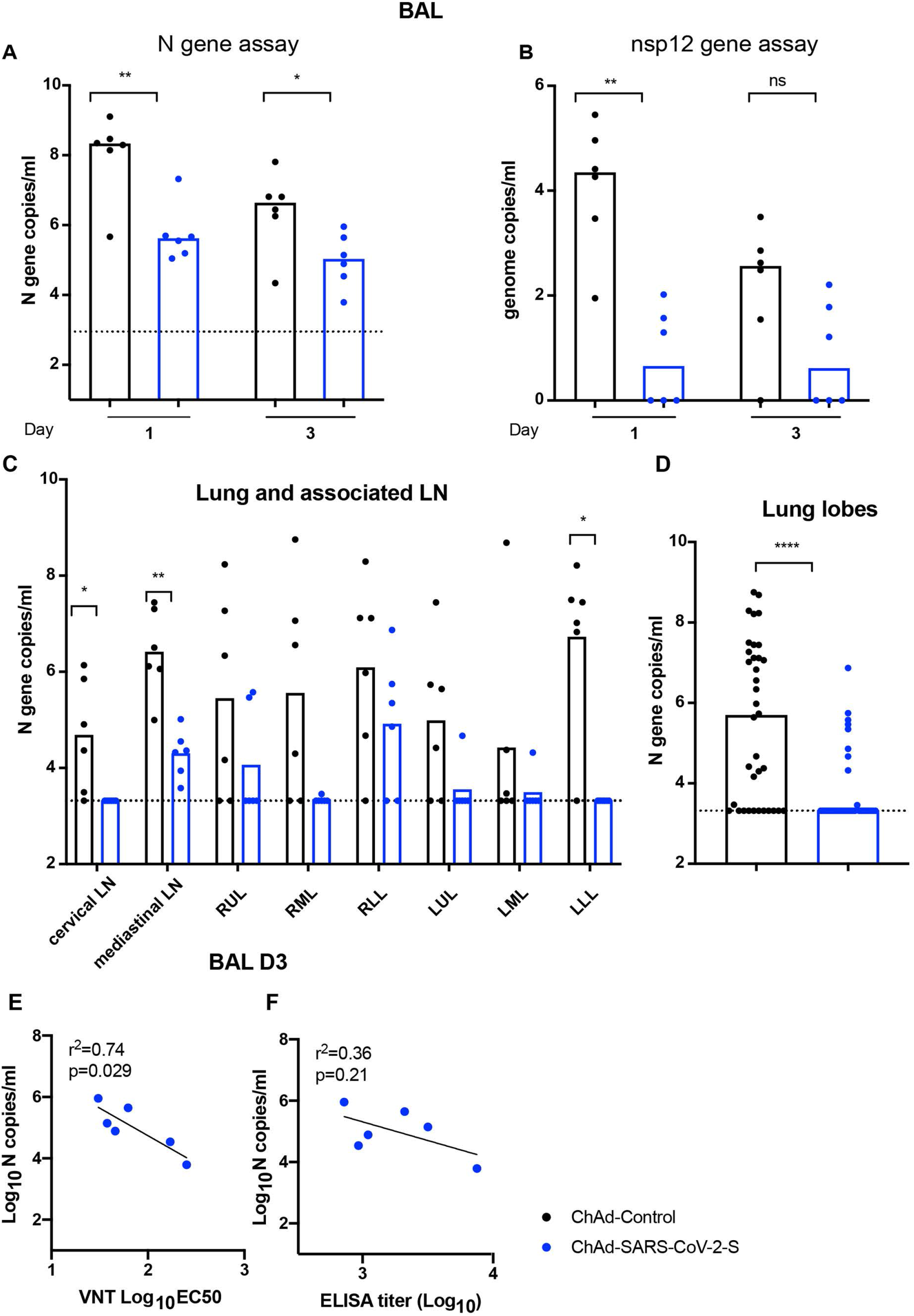
Viral RNA levels in lower respiratory tract of RM after SARS-CoV-2 challenge. RM (3-11 years old) immunized with ChAd-SARS-CoV-2-S or ChAd-Control were challenged with SARS-CoV-2 as described in **Figure 2. A-B.** BAL fluid was collected at day +1 and +3, and viral subgenomic (**A**) and genomic (**B**) RNA levels were assessed in BAL fluids by RT-qPCR) (n = 6, Mann-Whitney test: * *P* < 0.05; ** *P* < 0.01). **C-D.** Tissue samples from different lung lobes, mediastinal LN, and cervical LN were collected at day +7 after challenge. Subgenomic viral RNA levels were assessed by RT-qPCR (**C**). Viral RNA levels of the combined lung lobe specimens from ChAd-Control or ChAd-SARS-CoV-2-S immunized RM are shown (**D**) (n = 6, Mann-Whitney test: * *P* < 0.05; **** *P* < 0.0001). **E-F.** Correlation analysis of serum neutralizing antibody titers (**E**) and anti-S antibody levels (**F**) in vaccinated RM at three weeks post-immunization with viral RNA levels in BAL at day 3 after challenge. The black lines represent the linear regression fit (n = 6, Spearman correlation test: *P* and *r^2^* values shown).

**Table 1.** Virus isolation from BAL fluid

| Vaccine | RM # | BAL Day 1 | BAL Day 3 |
| --- | --- | --- | --- |
| ChAd-SARS-COV-2-S | 1 | - | - |
|  | 2 | - | - |
|  | 3 | - | - |
|  | 4 | - | - |
|  | 5 | - | - |
|  | 6 | + | - |
| ChAd-Control | 7 | + | - |
|  | 8 | ++ | - |
|  | 9 | +++ | ++ |
|  | 10 | ++ | - |
|  | 11 | + | - |
|  | 12 | + | - |

- : negative, + : $<10^2$ TCID<sub>50</sub>/ml; TCID<sub>50</sub>, tissue culture infectious dose 50, ++ : $10^2$ - $10^4$ TCID<sub>50</sub>/ml, and +++ : $10^5$ TCID<sub>50</sub>/ml

At day +7, all animals were euthanized, and tissues were collected. Viral RNA was detected in the cervical lymph nodes (LN), mediastinal LN, and the lung tissues in the majority of ChAd-Control vaccinated animals. However, in ChAd-SARS-CoV-2-S immunized animals, lower, if any, viral RNA was detected (**Fig 3C**). Indeed, the viral RNA levels in the combined lung lobes from all ChAd-SARS-CoV-2-S immunized animals were substantially lower than that measured in ChAd-Control-immunized animals (**Fig 3D**). To begin to establish correlates of protection, the viral RNA levels in BAL fluid at day +3 were compared to the serum neutralizing or anti-S IgG titers obtained three weeks after immunization. We observed an inverse correlation between viral RNA levels in BAL fluid obtained three days after SARS-CoV-2 challenge and neutralizing antibody titers (**Fig 3E**). The neutralizing antibody levels correlated better than anti-S IgG levels (P = 0.029, R^2^ = 0.74 versus P = 0.21, R^2^ = 0.36, respectively) (**Fig 3E-F**). Thus, serum neutralizing antibody titers may serve as a correlate of protection for the ChAd-SARS-CoV-2-S vaccine.

### Pathological analysis of lungs from vaccinated RM

In this particular set of challenge experiments, infection in RMs was mild, and chest radiographs did not show evidence of frank consolidative pneumonia. ChAd-Control vaccinated RMs developed changes consistent with mild pulmonary disease (**Fig 4A**). In two animals, we observed marked interstitial pneumonia characterized by small foci of alveolar septae thickened by edema fluid and fibrin with evidence of macrophage and neutrophil infiltration. Adjacent alveoli contained small numbers of foamy pulmonary macrophages and rare neutrophils and were occasionally lined by small numbers of type II pneumocytes. Perivascular infiltrates with small numbers of lymphocytes forming perivascular cuffs was observed. Immunohistochemistry revealed that 4 of the 6 ChAd-Control RMs were positive for viral antigen that principally localized to type I pneumocytes (**Fig 4B**). Two ChAd-SARS-CoV-2-S vaccinated RMs also showed small microscopic pulmonary lesions that were similar to the ChAd-Control animals (**Fig 4A**). Notwithstanding these findings, none of the ChAd-SARS-CoV-2-S vaccinated RMs showed evidence of viral antigen staining in lung tissues as analyzed by immunohistochemistry (**Fig 4B**).

**Figure 4.**
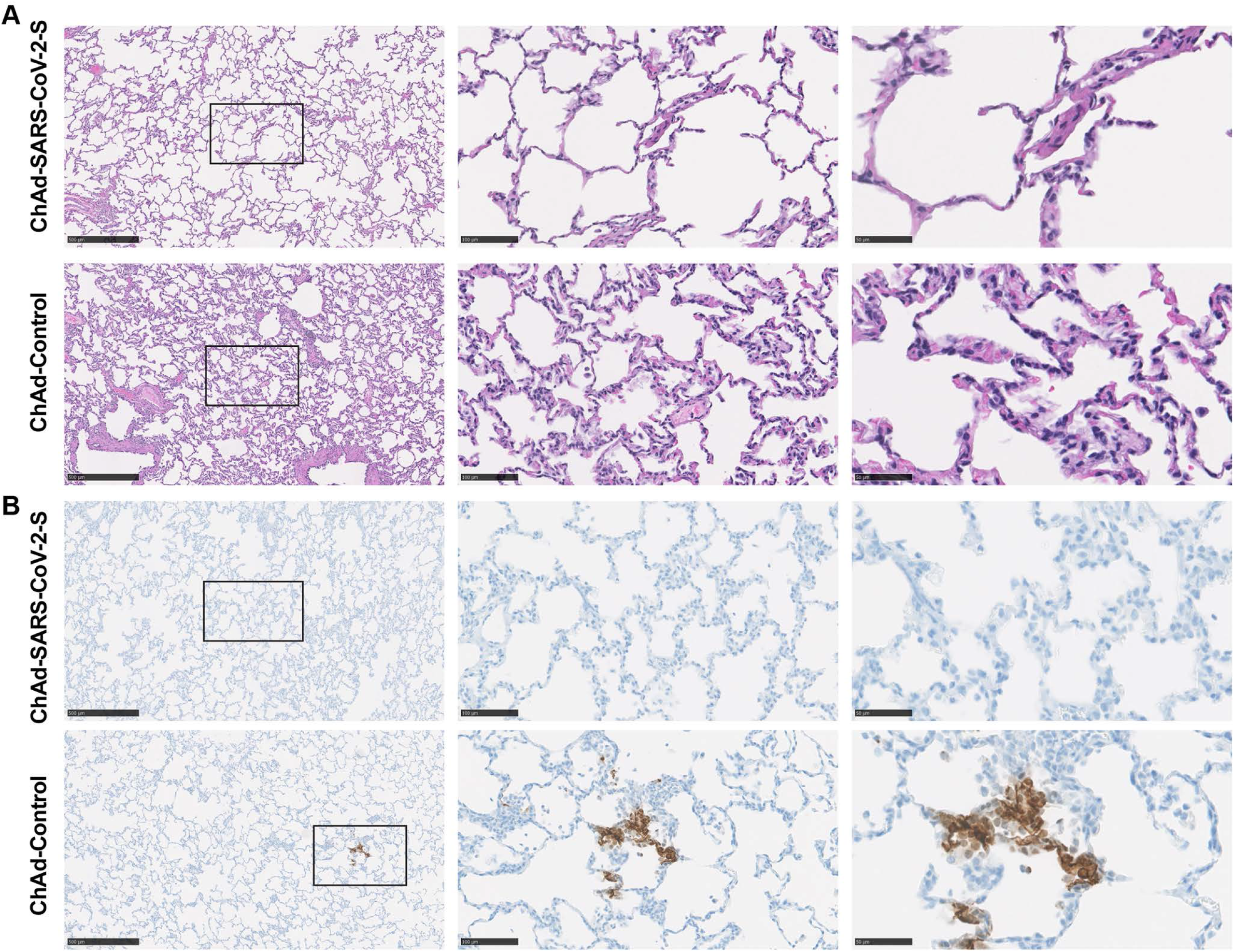
Pathological analysis of lungs of vaccinated RMs. RMs were immunized with ChAd-control and ChAd-SARS-CoV-2-S and challenged following the scheme described in **Fig 2**. Lungs were harvested at 7 dpi. **A**. Sections were stained with hematoxylin and eosin and imaged. Each image is representative of a group of 6 RMs. **B.** SARS-CoV-2 antigen was detected in lung sections from RMs for conditions described in (**A**). Images show low- (left; scale bars, 500 μm), medium- (middle; scale bars, 100 μm), and high-power magnification (right; scale bars, 50 μm). Representative images from n = 6 RMs per group.

## DISCUSSION

In this study, we show that in RM, a single intranasal immunization of ChAd-SARS-CoV-2 confers protection in both the upper and lower airways against challenge with a high dose of SARS-CoV-2. These results are consistent with recent studies showing protection against ChAd-SARS-CoV-2 after a similar immunization strategy in mice expressing hACE2 receptors (Hassan et al., 2020) and in hamsters (Bricker et al., 2020). Within three weeks of intranasal vaccination, we detected S protein-specific T cell responses in peripheral blood as well as anti-S IgG and IgA, anti-RBD, and neutralizing antibodies in serum. Immunization with ChAd-SARS-CoV-2 compared to the control ChAd-Control vaccine resulted in more rapid clearance of SARS-CoV-2 RNA from nasal samples, decreased levels of viral RNA in BAL fluid, and lower levels of viral RNA in the homogenates from different regions of the lungs.

Our experiments showing protection against SARS-CoV-2 infection and pathogenesis in RM are consistent with other studies in nonhuman primates using similar or distinct vaccine platforms. These include: (a) a three-dose intramuscular vaccination regimen with whole inactivated SARS-CoV-2, which protected RM from SARS-CoV-2-induced pneumonia (Gao et al., 2020); (b) a two-dose intramuscular vaccination regimen with a DNA vaccine encoding the S protein, which reduced viral RNA levels in BAL fluid and nasal swabs (Yu et al., 2020); (c) a priming or two-dose intramuscular immunization with ChAdOx1 nCoV-19, a related chimpanzee adenoviral vector, resulted in reduced viral loads in BAL fluid and lung tissues without protection against upper airway infection (van Doremalen et al., 2020); (d) intramuscular immunization with a single dose human Ad26 adenoviral vector provided near-complete protection in BAL fluid and nasal swabs after SARS-CoV-2 challenge (Mercado et al., 2020) although viral RNA levels in lung tissues were not measured; and (e) one month after intranasal or intramuscular vaccination with a human Ad5 adenoviral vector, RM were protected against SARS-CoV-2 challenge as evidenced by decreased viral RNA levels in oropharyngeal swabs and lung tissues (Feng et al., 2020). While all of these studies show protection in the RM model, it nonetheless is challenging to compare them directly because of the following disparities: RMs of different origin, different immunization and challenge protocols, different viral stocks and inoculation doses, different clinical and pathological scoring systems, separate assays for readout of infection, and different study locations.

The serum neutralization titers at 21 days with a single intranasal dose of ChAd-SARS-CoV-2 were similar to those obtained with other one or two-dose intramuscular vaccine platforms in RM. However, these levels are lower than what we observed in BALB/c or C57BL/6 mice immunized with a single dose of ChAd-SARS-CoV-2 (Hassan et al., 2020). The basis for this discrepancy remains uncertain, although higher doses on a viral particle/kg were used in mice. As reported recently by others, the relatively modest levels of immune response in RM after intranasal vaccination of adenoviral vectors still conferred effective protection against SARS-CoV-2 (Feng et al., 2020), and our data show that serum neutralizing antibody levels are a reasonable correlate of protection. These results are consistent with a recent study establishing serum antibody as a correlate of protection against SARS-CoV-2 infection in RMs (McMahan et al., 2020). In comparison, others have used anti-RBD responses in convalescent plasma from humans to predict protective immunity (Salazar et al., 2020). Future vaccine dose-response analysis in RM could provide further insight into the correlates of protection and potential ways to enhance the humoral response against SARS-CoV-2. Alternatively, a homologous intranasal or intramuscular booster dose (Richardson et al., 2013) with ChAd-SARS-CoV-2 or a heterologous adenoviral vector could augment immune responses and protection.

Vaccines for COVID-19 should protect against pneumonia and death and curtail transmission in the population. Our preclinical data in mice (Hassan et al., 2020), hamsters (Bricker et al., 2020), and RM suggest that ChAd-SARS-CoV-2 might accomplish these goals given its ability to reduce levels of subgenomic SARS-CoV-2 RNA and infectious virus in the BAL fluid and lungs and diminish infection at nasopharyngeal portal sites. We challenged RM with a high dose of virus (1 x 10^6^ TCID_50_) using two routes (intranasal and intrabronchial) to test the efficacy of the vaccine. As this likely does not reflect the exposure pattern in humans, the ChAd-SARS-CoV-2 vaccine may show better efficacy. Indeed, the direct intrabronchial inoculation of SARS-CoV-2 used in our challenge model, which is necessary to establish consistent pulmonary infection (Chandrashekar et al., 2020; Muñoz-Fontela et al., 2020), likely bypassed the nasal and pharyngeal mucosal immune barrier, and limited the protective impact of the intranasal vaccine.

In contrast to results with SARS-CoV vaccines or antibodies (Bolles et al., 2011; Liu et al., 2019; Weingartl et al., 2004), we did not observe enhanced infection, immunopathology, or enhanced disease in animals immunized with the ChAd-SARS-CoV-2 vaccine. In the ChAd-SARS-CoV-2 vaccinated and challenged RM, we did not observe worsening in clinical signs or enhanced replication compared to controls at any time point in the study, Moreover, the pathological analysis did not show evidence of the enhanced immune cell infiltration or alveolar damage seen in pulmonary tissues of vaccinated, SARS-CoV challenged RMs.

We acknowledge several limitations in this study: (a) we did not perform direct immunogenicity and efficacy comparisons with intramuscular delivery of ChAd-SARS-CoV-2; (b) we did not assess durability of immune responses. Future longitudinal studies must be conducted to monitor immune responses over time after intranasal vaccination with ChAd-SARS-CoV-2-S to establish durability; and (c) even with a high dose and invasive route of challenge, many of the control vaccinated animals did not develop severe lung pathology or disease, which limited our ability to conclude protection against disease in this model.

In summary, our studies establish that immunization of primates with ChAd-SARS-CoV-2-S induces neutralizing antibody and protective immune responses in both the upper and lower respiratory tract. While several vaccine candidates (mRNA, inactivated, viral-vectored) are in advanced phases of human clinical trials or recently granted EUAs for immunization of humans, their efficacy in curtailing transmission remains to be established. Based on preclinical data in multiple animal models, we suggest that intranasal delivery of ChAd-SARS-CoV-2-S or possibly other viral-vectored or subunit-based vaccines is a promising platform for preventing SARS-CoV-2 infection, disease, and transmission, and warrants further evaluation in humans.

## ACKNOWLEDGEMENTS

The authors thank the staff of the Rocky Mountain Veterinary Branch, NIAID, NIH for animal care and veterinary services. This study was supported by NIH contracts and grants (R01 AI157155, 75N93019C00062, HHSN272201400018C) as well as the Intramural Research Program of NIAID, NIH.

## AUTHOR CONTRIBUTIONS

A.O.H. generated the vaccines, performed ELISA assays, and analyzed the data. J.B.C. performed neutralization assays. H.Z. and D.H.F. designed and produced the recombinant S and RBD proteins. T.L.T.H. performed the T cell assays. F.F., A.O., J.L., and P.W.H. performed and evaluated the clinical exams, sample collection, hematology and blood chemistry and clinical scoring. F.F., K.M.W., T.L.T.H. and D.P.S performed the necropsies and organ harvest. D.P.S. performed and evaluated the histopathology. A.O.H., F.F., J.C. and K.M.W. performed and evaluated the virological assays. H.F. and M.S.D. designed experiments and secured funding. D.T.C. provided key vaccine reagents. A.O.H., H.F., and M.S.D. wrote the initial draft, with the other authors providing editorial comments.

---

**Figure S1. Absence of anti-vector antibodies in RMs**. Serum samples were collected from RMs on day −28 prior to ChAd immunization and evaluated for neutralization activity against ChAd using a FRNT. Each symbol represents a single animal; each point represents data from two technical repeats.

## STAR METHODS

### RESOURCE AVAILABLITY

#### Lead Contact

Further information and requests for resources and reagents should be directed to and will be fulfilled by the Lead Contact, Michael S. Diamond.

#### Materials Availability

All requests for resources and reagents should be directed to and will be fulfilled by the Lead Contact author. This includes viruses, vaccines, and proteins. All reagents will be made available on request after completion of a Materials Transfer Agreement.

#### Data and code availability

All data supporting the findings of this study are available within the paper and are available from the corresponding author upon request.

### EXPERIMENTAL MODEL AND SUBJECT DETAILS

#### Viruses and cells

Vero E6 (CRL-1586, American Type Culture Collection (ATCC), and HEK293 cells were cultured at 37°C in Dulbecco’s Modified Eagle medium (DMEM) supplemented with 10% fetal bovine serum (FBS), 10 mM HEPES pH 7.3, 1 mM sodium pyruvate, 1X non-essential amino acids, and 100 U/ml of penicillin-streptomycin. SARS-CoV-2 strain 2019 n-CoV/USA_WA1/2020 was kindly provided by the Centers for Disease Control and Prevention as a passage 3 stock. The virus was propagated once more in Vero E6 cells in DMEM (Sigma) supplemented with 2% fetal bovine serum (Gibco), 1 mM L-glutamine (Gibco), 50 U/ml penicillin and 50 μg/ml of streptomycin (Gibco). The virus stock used was passage 4, free of mycoplasma contamination, and identical to the initial deposited Genbank sequence (MN985325.1).

#### RM experiments

Twelve healthy rhesus macaques (*Macaca mulatta*; Indian origin, between 3 and 11 years of age and 4 – 10 kg in weight) were divided randomly into two groups of 6 animals (3 females and 3 males). All macaques were immunized 28 days prior to challenge by the intranasal route with a total dose of 10^11^ viral particles of ChAd-Control or ChAd-SARS-CoV-2-S in a total volume of 1 mL (0.5 mL into each nostril). Challenge was performed with a total dose of 1 x10^6^ TCID_50_ of SARS-CoV-2 equally split between the two routes in a total volume of 5 mL (4 mL intrabronchial; 1 mL intranasal). Animals were monitored at least twice daily throughout the study using an established scoring sheet (Brining et al., 2010; Munster et al., 2020). Examinations were performed on days −28, −27, −25, −21, −14, −7, 0, 1, 3, 5 and 7 (euthanasia) and included clinical evaluation, thoracic radiographs, venous blood draw, and swabs. Bronchoalveolar lavage (BAL) was performed on day 1 and 3 as described (Singletary et al., 2008). The study endpoint was day 7. Following euthanasia, necropsies were performed on all animals, and organs were harvested for virology, immunology and pathology. Gross lung lesions were scored by a board-certified veterinary pathologist blinded to the group assignment. Animals were sedated with either ketamine (10-12 mg/Kg) or Telazol (3.5-5 mg/Kg) by intramuscular injection for all clinical examinations and procedures. Isoflurane in oxygen was administered via face mask as needed to maintain appropriate levels of sedation and anesthesia.

### METHOD DETAILS

#### Biosafety and ethics

Work with infectious SARS-CoV-2 was approved by the Institutional Biosafety Committee (IBC) and performed in high biocontainment at Rocky Mountain Laboratories (RML), NIAID, NIH. Sample removal from high biocontainment followed IBC-approved Standard Operating Protocols. Animal work was approved by the RML Animal Care and Use Committee and performed by certified staff in an Association for Assessment and Accreditation of Laboratory Animal Care International accredited facility. Work followed the institution’s guidelines for animal use, the guidelines and basic principles in the NIH Guide for the Care and Use of Laboratory Animals, the Animal Welfare Act, United States Department of Agriculture and the United States Public Health Service Policy on Humane Care and Use of Laboratory Animals. Nonhuman primates were single housed in adjacent primate cages allowing social interactions, in a climate-controlled room with a fixed light-dark cycle (12-hr light/12-hr dark). They were provided with commercial monkey chow, treats, and fruit twice daily with water *ad libitum*. Environmental enrichment consisted of a variety of human interaction, manipulanda, commercial toys, videos, and music.

#### Chimpanzee adenovirus vectors

The replication-incompetent ChAd-SARS-CoV-2-S and ChAd-Control vectors were previously described (Hassan et al., 2020). Vaccines were scaled up in 293 cells and purified by CsCl density-gradient ultracentrifugation. Viral particle concentration in each vector preparation was determined by spectrophotometry at 260 nm as described (Maizel et al., 1968).

#### Neutralization assay

Serum samples were diluted serially and incubated with 10^2^ FFU of SARS-CoV-2 for 1 h at 37°C. The virus-serum mixtures were added to Vero E6 cell monolayers in 96-well plates and incubated for 1 h at 37°C. Subsequently, cells were overlaid with 1% (w/v) methylcellulose in MEM supplemented with 2% FBS. Plates were incubated for 30 h then fixed using 4% PFA in PBS for 1 h at room temperature. After washing, cells were sequentially incubated with murine anti-SARS-CoV-2 S mAb (L. VanBlargan and M. Diamond, unpublished results) and a HRP-conjugated goat anti-mouse IgG (Sigma) in PBS supplemented with 0.1% (w/v) saponin (Sigma) and 0.1% BSA. TrueBlue peroxidase substrate (KPL) was used to develop the plates followed by counting the foci on a BioSpot analyzer (Cellular Technology Limited).

#### ELISA

Purified antigens (S or RBD) were coated onto 96-well Maxisorp clear plates at 2 μg/mL in 50 mM Na_2_CO_3_ pH 9.6 (70 μL) overnight at 4°C. Coating buffers were aspirated, and wells were blocked with 200 μL of PBS + 0.05% Tween-20 + 5 % BSA (Blocking buffer, PBSTBA) overnight at 4°C. Serum samples were diluted in PBSTBA in a separate 96-well polypropylene plate. The plates then were washed thrice with 1X PBS + 0.05% Tween-20 (PBST) followed by addition of 50 μL of respective serum dilutions. Sera were incubated in the ELISA plates for at least 1 h at room temperature. Plates were again washed thrice in PBST followed by addition of 50 μL of 1:1000 goat anti-RM IgG(H+L)-HRP (Southern Biotech Cat. # 6200-05) in PBSTBA. Plates were incubated at room temperature for 1 h, washed thrice in PBST, and 100 μL of 1-Step Ultra TMB-ELISA was added (ThermoFisher Cat. #34028). Reactions were stopped with 50 μL of 2 M sulfuric acid. Optical density (450 nm) measurements were determined using a microplate reader (Bio-Rad).

#### Simian Ad neutralization assays

Serum samples were collected from RMs one day prior to ChAd immunizations. Sera were serially diluted prior to incubation with 10^2^ FFU of ChAd-SARS-CoV-2-S for 1 h at 37°C. The virus-serum mixtures were added to Vero cell monolayers in 96-well plates and incubated for 1 h at 37°C. Cells were overlaid with 1% (w/v) methylcellulose in MEM supplemented with 5% FBS. Plates were incubated at 37°C for 48 h before fixation with 4% PFA in PBS for 20 min at room temperature. Subsequently, plates were washed with PBS and incubated overnight at 4°C with murine anti-SARS-CoV-2 S mAb (L. VanBlargan and M. Diamond, unpublished results) diluted in permeabilization buffer (PBS supplemented with 0.1% (w/v) saponin and 0.1% BSA). Plates were washed again and incubated with anti-mouse-HRP (1:500; Sigma Cat. # A9044) in permeabilization buffer for 1 h at room temperature. After a final wash series, plates were developed using TrueBlue peroxidase substrate (KPL) and foci were counted on a BioSpot analyzer (Cellular Technology Limited).

#### T cell assay

Peripheral blood mononuclear cells (PBMCs) were prepared by centrifugation on a Ficoll gradient. Cells (3.0 x 10^6^ PBMCs) from each RM were seeded into duplicate wells in a 96-well flat-bottom plate which pre-coated with human IFN-γ-capturing antibody (3.0 × 10^5^ cells/well-Human IFN-γ Single-Color ELISPOT-ImmunoSpot). PBMCs were stimulated with SARS-CoV-2 spike protein peptide pools, at a final concentration of 2 μg/mL per peptide, for 24 h in a humidified incubator with 5% CO2 at 37 °C. IFN-γ spots were developed according to the manufacturer’s protocol and counted by CTL 328 ImmunoSpot^®^ Analyzer and ImmunoSpot^®^ Software.

#### Measurement of viral burden

SARS-CoV-2 infected animals were euthanized at day +7 after SARS-CoV-2 challenge, and tissues were collected. Tissues (up to 30 mg) were homogenized in RLT buffer. RNA was extracted using RNeasy kit (Qiagen) according to manufacturer’s instructions. RNA was extracted from BAL fluid and nasal swabs using the QiaAmp Viral RNA kit (Qiagen) according to manufacturer’s instructions. SARS-CoV-2 RNA levels were measured by one-step TaqMan RT-qPCR assay. SARS-CoV-2 nucleocapsid (N) or nsp12 specific primers and probe sets were used: (N: F primer: ATGCTGCAATCGTGCTACAA; R primer: GACTGCCGCCTCTGCTC; probe: /56-FAM/TCAAGGAAC/ZEN/AACATTGCCAA/3IABkFQ/ and nsp12: F primer: GTGARATGGTCATGTGTGGCGG; R primer: CARATGTTAAASACACTATTAGCATA; probe 1: 6-FAM/CCAGGTGGWACRTCATCMGGTGATGC; probe 2: 6-FAM/CAGGTGGAACCTCATCAGGAGATGC. Viral RNA was expressed as N or nsp12 gene copy numbers per g of tissues or per ml of fluid.

Virus isolation was performed on BAL liquid and homogenized lung tissues (approximately 30 mg in 1 mL DMEM using a TissueLyser (Qiagen)) and inoculating Vero E6 cells in a 24 well plate with 250 μL of cleared and a 1:10 dilution of the homogenate. One hour after inoculation of cells, the supernatant was removed and replaced with 500 μL DMEM (Sigma-Aldrich) supplemented with 2% fetal bovine serum, 1 mM L-glutamine, 50 U/mL penicillin and 50 μg/mL of streptomycin. Six days after inoculation, cytopathogenic effect was scored, and the TCID_50_ was calculated.

#### Histopathology and immunohistochemistry

Histopathology and immunohistochemistry were performed on RM tissues. After fixation for a minimum of 7 days in 10% neutral-buffered formalin and embedding in paraffin, tissue sections were stained with hematoxylin and eosin (H & E). Tissues were placed in cassettes and processed with a Sakura VIP-6 Tissue Tek on a 12-hour automated schedule, using a graded series of ethanol, xylene, and ParaPlast Extra. Embedded tissues are sectioned at 5 μm and dried overnight at 42°C prior to staining. Specific anti-SARS-CoV-2 immunoreactivity was detected using GenScript U864YFA140-4/CB2093 NP-1 at a 1:1,000 dilution. The secondary antibody used was an antirabbit IgG polymer from Vector Laboratories ImPress VR. Tissues were processed for immunohistochemistry using the Discovery Ultra automated processor (Ventana Medical Systems) with a ChromoMap DAB kit (Roche Tissue Diagnostics).

### QUANTIFICATION AND STATISTICAL ANALYSIS

Statistical significance was assigned when *P* values were < 0.05 using Prism Version 8 (GraphPad). Tests, number of animals (n), median values, and statistical comparison groups are indicated in the Figure legends.

## REFERENCES

Baden, L.R., El Sahly, H.M., Essink, B., Kotloff, K., Frey, S., Novak, R., Diemert, D., Spector, S.A., Rouphael, N., Creech, C.B., et al. (2020). Efficacy and Safety of the mRNA-1273 SARS-CoV-2 Vaccine. N Engl J Med.

Barnes, C.O., Jette, C.A., Abernathy, M.E., Dam, K.A., Esswein, S.R., Gristick, H.B., Malyutin, A.G., Sharaf, N.G., Huey-Tubman, K.E., Lee, Y.E., et al. (2020). SARS-CoV-2 neutralizing antibody structures inform therapeutic strategies. Nature 588, 682–687.

Bolles, M., Deming, D., Long, K., Agnihothram, S., Whitmore, A., Ferris, M., Funkhouser, W., Gralinski, L., Totura, A., Heise, M., et al. (2011). A double-inactivated severe acute respiratory syndrome coronavirus vaccine provides incomplete protection in mice and induces increased eosinophilic proinflammatory pulmonary response upon challenge. J Virol 85, 12201–12215.

Bricker, T.L., Darling, T.L., Hassan, A.O., Harastani, H.H., Soung, A., Jiang, X., Dai, Y.N., Zhao, H., Adams, L.J., Holtzman, M.J., et al. (2020). A single intranasal or intramuscular immunization with chimpanzee adenovirus vectored SARS-CoV-2 vaccine protects against pneumonia in hamsters. bioRxiv.

Brining, D.L., Mattoon, J.S., Kercher, L., LaCasse, R.A., Safronetz, D., Feldmann, H., and Parnell, M.J. (2010). Thoracic radiography as a refinement methodology for the study of H1N1 influenza in cynomologus macaques (Macaca fascicularis). Comparative medicine 60, 389–395.

Burton, D.R., and Walker, L.M. (2020). Rational Vaccine Design in the Time of COVID-19. Cell Host Microbe 27, 695–698.

Cao, L., Goreshnik, I., Coventry, B., Case, J.B., Miller, L., Kozodoy, L., Chen, R.E., Carter, L., Walls, A.C., Park, Y.J., et al. (2020a). De novo design of picomolar SARS-CoV-2 miniprotein inhibitors. Science 370, 426–431.

Cao, Y., Su, B., Guo, X., Sun, W., Deng, Y., Bao, L., Zhu, Q., Zhang, X., Zheng, Y., Geng, C., et al. (2020b). Potent neutralizing antibodies against SARS-CoV-2 identified by high-throughput single-cell sequencing of convalescent patients’ B cells. Cell 182, 73–84.

Case, J.B., Rothlauf, P.W., Chen, R.E., Liu, Z., Zhao, H., Kim, A., S., Bloyet, L.M., Zeng, Q., Tahan, S., Droit, L., et al. (2020). Neutralizing antibody and soluble ACE2 inhibition of a replication-competent VSV-SARS-CoV-2 and a clinical isolate of SARS-CoV-2. Cell Host and Microbe 28, 475–485.

Chandrashekar, A., Liu, J., Martinot, A.J., McMahan, K., Mercado, N.B., Peter, L., Tostanoski, L.H., Yu, J., Maliga, Z., Nekorchuk, M., et al. (2020). SARS-CoV-2 infection protects against rechallenge in rhesus macaques. Science 369, 812–817.

Cheung, E.W., Zachariah, P., Gorelik, M., Boneparth, A., Kernie, S.G., Orange, J.S., and Milner, J.D. (2020). Multisystem Inflammatory Syndrome Related to COVID-19 in Previously Healthy Children and Adolescents in New York City. JAMA 324, 294–296.

Feng, L., Wang, Q., Shan, C., Yang, C., Feng, Y., Wu, J., Liu, X., Zhou, Y., Jiang, R., Hu, P., et al. (2020). An adenovirus-vectored COVID-19 vaccine confers protection from SARS-COV-2 challenge in rhesus macaques. Nat Commun 11, 4207.

Gao, Q., Bao, L., Mao, H., Wang, L., Xu, K., Yang, M., Li, Y., Zhu, L., Wang, N., Lv, Z., et al. (2020). Rapid development of an inactivated vaccine candidate for SARS-CoV-2. Science 369, 77–81.

Graham, B.S. (2020). Rapid COVID-19 vaccine development. Science 368, 945–946.

Guan, W.J., Ni, Z.Y., Hu, Y., Liang, W.H., Ou, C.Q., He, J.X., Liu, L., Shan, H., Lei, C.L., Hui, D. S.C., et al. (2020). Clinical Characteristics of Coronavirus Disease 2019 in China. N Engl J Med.

Hassan, A.O., Kafai, N.M., Dmitriev, I.P., Fox, J.M., Smith, B.K., Harvey, I.B., Chen, R.E., Winkler, E. S., Wessel, A.W., Case, J.B., et al. (2020). A Single-Dose Intranasal ChAd Vaccine Protects Upper and Lower Respiratory Tracts against SARS-CoV-2. Cell 183, 169–184.e113.

Hoffmann, M., Kleine-Weber, H., Schroeder, S., Kruger, N., Herrler, T., Erichsen, S., Schiergens, T.S., Herrler, G., Wu, N.H., Nitsche, A., et al. (2020). SARS-CoV-2 Cell Entry Depends on ACE2 and TMPRSS2 and Is Blocked by a Clinically Proven Protease Inhibitor. Cell.

Letko, M., Marzi, A., and Munster, V. (2020). Functional assessment of cell entry and receptor usage for SARS-CoV-2 and other lineage B betacoronaviruses. Nature microbiology 5, 562–569.

Liu, L., Wei, Q., Lin, Q., Fang, J., Wang, H., Kwok, H., Tang, H., Nishiura, K., Peng, J., Tan, Z., et al. (2019). Anti-spike IgG causes severe acute lung injury by skewing macrophage responses during acute SARS-CoV infection. JCI insight 4.

Maizel, J.V., Jr., White, D.O., and Scharff, M.D. (1968). The polypeptides of adenovirus. I. Evidence for multiple protein components in the virion and a comparison of types 2, 7A, and 12. Virology 36, 115–125.

Mao, R., Qiu, Y., He, J.S., Tan, J.Y., Li, X.H., Liang, J., Shen, J., Zhu, L.R., Chen, Y., Iacucci, M., et al. (2020). Manifestations and prognosis of gastrointestinal and liver involvement in patients with COVID-19: a systematic review and meta-analysis. The lancet Gastroenterology & hepatology.

McMahan, K., Yu, J., Mercado, N.B., Loos, C., Tostanoski, L.H., Chandrashekar, A., Liu, J., Peter, L., Atyeo, C., Zhu, A., et al. (2020). Correlates of protection against SARS-CoV-2 in rhesus macaques. Nature.

Mercado, N.B., Zahn, R., Wegmann, F., Loos, C., Chandrashekar, A., Yu, J., Liu, J., Peter, L., McMahan, K., Tostanoski, L.H., et al. (2020). Single-shot Ad26 vaccine protects against SARS-CoV-2 in rhesus macaques. Nature 586, 583–588.

Muñoz-Fontela, C., Dowling, W.E., Funnell, S.G.P., Gsell, P.S., Riveros-Balta, A.X., Albrecht, R.A., Andersen, H., Baric, R.S., Carroll, M.W., Cavaleri, M., et al. (2020). Animal models for COVID-19. Nature 586, 509–515.

Munster, V.J., Feldmann, F., Williamson, B.N., van Doremalen, N., Pérez-Pérez, L., Schulz, J., Meade-White, K., Okumura, A., Callison, J., Brumbaugh, B., et al. (2020). Respiratory disease in rhesus macaques inoculated with SARS-CoV-2. Nature 585, 268–272.

Pinto, D., Park, Y.J., Beltramello, M., Walls, A.C., Tortorici, M.A., Bianchi, S., Jaconi, S., Culap, K., Zatta, F., De Marco, A., et al. (2020). Cross-neutralization of SARS-CoV-2 by a human monoclonal SARS-CoV antibody. Nature 583, 290–295.

Polack, F.P., Thomas, S.J., Kitchin, N., Absalon, J., Gurtman, A., Lockhart, S., Perez, J.L., Pérez Marc, G., Moreira, E.D., Zerbini, C., et al. (2020). Safety and Efficacy of the BNT162b2 mRNA Covid-19 Vaccine. N Engl J Med 383, 2603–2615.

Richardson, J.S., Pillet, S., Bello, A.J., and Kobinger, G.P. (2013). Airway delivery of an adenovirus-based Ebola virus vaccine bypasses existing immunity to homologous adenovirus in nonhuman primates. J Virol 87, 3668–3677.

Salazar, E., Kuchipudi, S.V., Christensen, P.A., Eagar, T., Yi, X., Zhao, P., Jin, Z., Long, S.W., Olsen, R.J., Chen, J., et al. (2020). Convalescent plasma anti-SARS-CoV-2 spike protein ectodomain and receptor binding domain IgG correlate with virus neutralization. J Clin Invest 130, 6728–6738.

Singletary, M.L., Phillippi-Falkenstein, K.M., Scanlon, E., Bohm, R.P., Jr., Veazey, R.S., and Gill, A.F. (2008). Modification of a common BAL technique to enhance sample diagnostic value. Journal of the American Association for Laboratory Animal Science: JAALAS 47, 47–51.

Tortorici, M.A., Beltramello, M., Lempp, F.A., Pinto, D., Dang, H.V., Rosen, L.E., McCallum, M., Bowen, J., Minola, A., Jaconi, S., et al. (2020). Ultrapotent human antibodies protect against SARS-CoV-2 challenge via multiple mechanisms. Science 370, 950–957.

van Doremalen, N., Lambe, T., Spencer, A., Belij-Rammerstorfer, S., Purushotham, J.N., Port, J.R., Avanzato, V.A., Bushmaker, T., Flaxman, A., Ulaszewska, M., et al. (2020). ChAdOx1 nCoV-19 vaccine prevents SARS-CoV-2 pneumonia in rhesus macaques. Nature 586, 578–582.

Wang, H., Zhang, Y., Huang, B., Deng, W., Quan, Y., Wang, W., Xu, W., Zhao, Y., Li, N., Zhang, J., et al. (2020). Development of an Inactivated Vaccine Candidate, BBIBP-CorV, with Potent Protection against SARS-CoV-2. Cell 182, 713–721.e719.

Weingartl, H., Czub, M., Czub, S., Neufeld, J., Marszal, P., Gren, J., Smith, G., Jones, S., Proulx, R., Deschambault, Y., et al. (2004). Immunization with modified vaccinia virus Ankara-based recombinant vaccine against severe acute respiratory syndrome is associated with enhanced hepatitis in ferrets. J Virol 78, 12672–12676.

Wichmann, D., Sperhake, J.P., Lütgehetmann, M., Steurer, S., Edler, C., Heinemann, A., Heinrich, F., Mushumba, H., Kniep, I., Schröder, A.S., et al. (2020). Autopsy Findings and Venous Thromboembolism in Patients With COVID-19: A Prospective Cohort Study. Ann Intern Med 173, 268–277.

Wrapp, D., Wang, N., Corbett, K.S., Goldsmith, J.A., Hsieh, C.L., Abiona, O., Graham, B.S., and McLellan, J.S. (2020). Cryo-EM structure of the 2019-nCoV spike in the prefusion conformation. Science 367, 1260–1263.

Yu, J., Tostanoski, L.H., Peter, L., Mercado, N.B., McMahan, K., Mahrokhian, S.H., Nkolola, J.P., Liu, J., Li, Z., Chandrashekar, A., et al. (2020). DNA vaccine protection against SARS-CoV-2 in rhesus macaques. Science 369, 806–811.

Zhou, F., Yu, T., Du, R., Fan, G., Liu, Y., Liu, Z., Xiang, J., Wang, Y., Song, B., Gu, X., et al. (2020a). Clinical course and risk factors for mortality of adult inpatients with COVID-19 in Wuhan, China: a retrospective cohort study. Lancet 395, 1054–1062.

Zhou, P., Yang, X.L., Wang, X.G., Hu, B., Zhang, L., Zhang, W., Si, H.R., Zhu, Y., Li, B., Huang, C.L., et al. (2020b). A pneumonia outbreak associated with a new coronavirus of probable bat origin. Nature 579, 270–273.

Zost, S.J., Gilchuk, P., Chen, R.E., Case, J.B., Reidy, J.X., Trivette, A., Nargi, R.S., Sutton, R.E., Suryadevara, N., Chen, E.C., et al. (2020). Rapid isolation and profiling of a diverse panel of human monoclonal antibodies targeting the SARS-CoV-2 spike protein. Nat Med 26, 1422–1427.

