## Supplemental Figure 1 and Table 1 for "A single intranasal dose of chimpanzee adenovirus-vectored vaccine protects against SARS-CoV-2 infection in rhesus macaques"

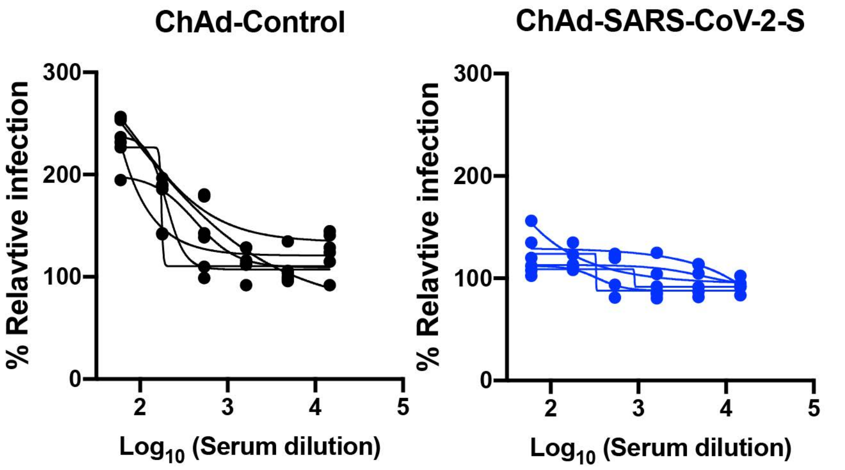
ChAd-Control
ChAd-SARS-CoV-2-S
% Relavtive infection
300
200
100
0
2
3
4
5
Log10 (Serum dilution)

**Table S1. Clinical scoring in ChAd vaccinated animals**

| Vaccine | RM # | Day 0-3 | Day 4-7 |
|---|---|---|---|
| ChAd-SARS-COV-2-S | 1 | Pale appearance, slow, deep irregular abdominal respirations<br>Score: 0/0/7/7 | Pale appearance, slow, deep irregular abdominal respirations<br>Score: 7/0/0/0 |
| | 2 | Normal<br>Score: 0/0/0/0 | Normal<br>Score: 0/0/0/0 |
| | 3 | Slightly ruffled fur, increased irregular respirations<br>Score: 0/2/7/7 | Slightly ruffled fur, irregular respirations<br>Score: 4/4/4/4 |
| | 4 | Reduced appetite<br>Score: 0/3/3/0 | normal<br>Score: 0/0/0/0 |
| | 5 | Slightly irregular abdominal respirations<br>Score:0/0/0/2 | Slightly irregular abdominal respirations<br>Score:2/0/2/2 |
| | 6 | Normal<br>Score: 0/3/0 | Slightly irregular abdominal respirations<br>Score: 2/0/0/0 |
| ChAd-Control | 7 | Normal<br>Score:0/0/0/0 | Slightly irregular abdominal respirations<br>Score: 0/2/0/2 |
| | 8 | Slightly ruffled fur, slightly irregular abdominal respirations<br>Score:0/2/4/4 | Slightly ruffled fur<br>Score: 2/0/0/0 |
| | 9 | Severely decreased appetite, ruffled fur, increased irregular abdominal respirations<br>Score:0/12/12/10 | Reduced appetite, increased irregular abdominal respirations<br>Score: 13/10/7/7 |
| | 10 | Ruffled fur, increased irregular abdominal respirations<br>Score: 0/5/5/7 | Ruffled fur, Increased irregular abdominal respirations<br>score: 10/7/10/10 |
| | 11 | Pale appearance, ruffled fur, increased irregular abdominal respirations<br>Score: 0/10/10/10 | Pale appearance, slow& tired, ruffled fur, increased irregular abdominal respirations, elevated temperature on day 5&7(39.9 & 39.7 °C<br>Score: 13/12/12/15 |
| | 12 | Pale appearance<br>Score: 0/0/0/0 | Pale appearance<br>Score: 0/0/0/0 |
